# Modeling the mutation and reversal of engineered underdominance gene drives

**DOI:** 10.1101/257253

**Authors:** Matthew P. Edgington, Luke S. Alphey

## Abstract

A range of gene drive systems have been proposed that are predicted to increase their frequency and that of associated desirable genetic material even if they confer a fitness cost on individuals carrying them. Engineered underdominance (UD) is such a system and, in one version, is based on the introduction of two independently segregating transgenic constructs each carrying a lethal gene, a suppressor for the lethal at the other locus and a desirable genetic “cargo”. Under this system individuals carrying at least one copy of each construct (or no copies of either) are viable whilst those that possess just one of the transgenic constructs are non-viable. Previous theoretical work has explored various properties of these systems, concluding that they should persist indefinitely in absence of resistance or mutation. Here we study a population genetics model of UD gene drive that relaxes past assumptions by allowing for loss-of-function mutations in each introduced gene. We demonstrate that mutations are likely to cause UD systems to break down, eventually resulting in the elimination of introduced transgenes. We then go on to investigate the potential of releasing “free suppressor” carrying individuals as a new method for reversing UD gene drives and compare this to the release of wild-types; the only previously proposed reversal strategy for UD. This reveals that while free suppressor carrying individuals may represent an inexpensive reversal strategy due to extremely small release requirements, they are not able to return a fully wild-type population as rapidly as the release of wild-types.

## 1 Introduction

Gene drive systems have gained much attention in recent years for their predicted ability to increase the frequency of desirable genetic material within a population. These systems have been proposed to have a number of important applications [1]. Our interest in these systems relates to their potential use in preventing the spread of mosquito-borne viruses such as dengue [2]. In this context, refractory genes have been developed that are capable of significantly reducing the ability of *Aedes aegypti* mosquitoes to transmit the virus [3]. However, the incorporation of these genes into the genome resulted in individuals of lower fitness than their wild-type counterparts [4]. It is thus necessary to develop gene drives that are capable of increasing the frequency of such desirable genes in spite of their expected fitness costs.

Engineered underdominance (UD) is one class of gene drive that has been proposed for driving desirable genetic traits into a population [5]. This technique is based on the introduction of two transgenic constructs (A and B) at unlinked genetic loci (see Figure 1). These constructs consist of three component genes, namely a lethal effector, a suppressor for the lethal at the other locus and a “cargo” gene conferring a desirable phenotype. Due to the cross suppressing transgenic constructs, individuals carrying one or more copies of either construct will be non-viable if they do not also possess at least one copy of the other construct. Individuals carrying one or more copies of both constructs are viable since each of the lethal effectors are inactivated by suppressors carried at the other locus. These effects combine to create a selection pressure for individuals to either carry both transgenic constructs or neither.

**Figure 1:**
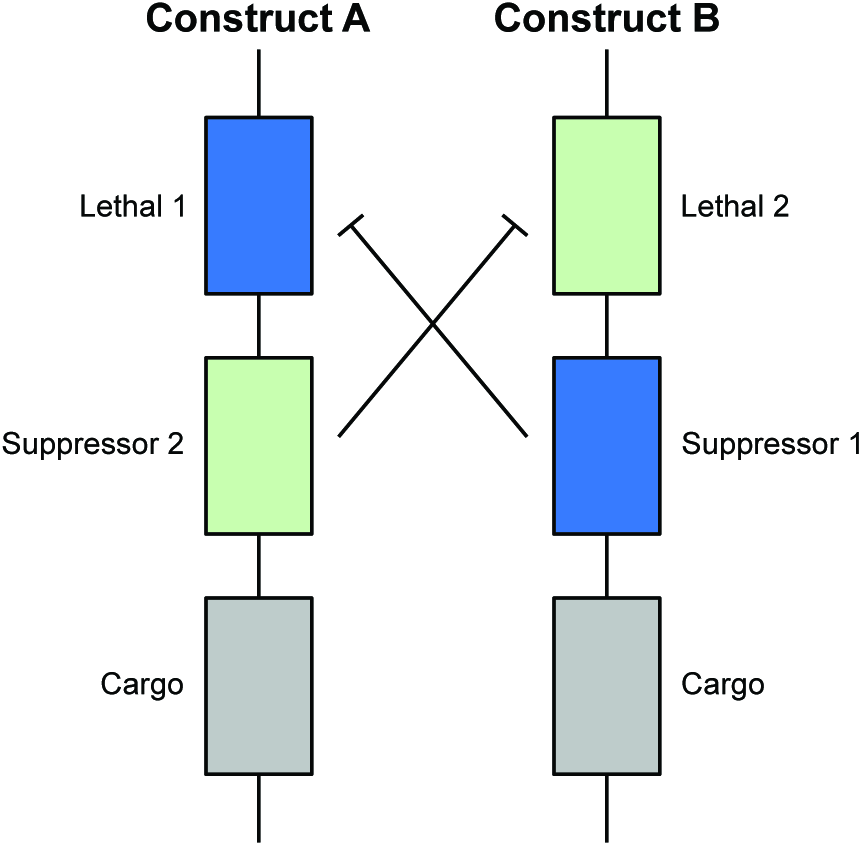
A schematic diagram of the engineered underdominance gene drive system. Each transgenic construct possesses three genes; a lethal, a suppressor for the lethal on the other construct and a desirable genetic cargo. Genotypes possessing one or more copies of both constructs (or none of either) are viable since lethals will be deactivated by the suppressor(s) on the other construct. Those genotypes carrying one or more copies of either construct but none of the other are non-viable since they have non-suppressed lethal genes.

Theoretical work on UD has demonstrated that these systems, when introduced above a threshold frequency, are capable of spreading desirable genes thus replacing wild populations with those carrying UD transgenic constructs [5-9]. Whilst the existence of this threshold frequency means that UD systems may be relatively expensive to deploy in the field relative to other classes of gene drive, it should restrict the invasion of transgenes into neighboring populations [10, 11].

Previous modeling work has mostly assumed a population genetics framework neglecting the possibility of mutations forming within the introduced transgenes [5-11]. Under this assumption it has been predicted that UD should persist indefinitely when introduced above the threshold frequency.

This threshold-dependent nature of UD systems has lead many to suggest that they may be reversed via the introduction of wild-type individuals [12, 13]. This should lower the transgene frequency to sub-threshold levels and result in the elimination of introduced transgenes. To our knowledge this is the only reversal strategy proposed for UD gene drives and has yet to be investigated in detail.

Here we extend upon results in the previous literature by formulating a population genetics model of UD that incorporates the effects of loss-of-function mutations in the introduced transgenes. In particular, we investigate the predicted dynamics of UD systems in the presence of a constant rate of mutation for each introduced transgene. We then go on to propose that the release of individuals carrying “free suppressors” could function as a genetics-based reversal strategy and compare this to the introduction of wild-types.

## 2 Mathematical Modeling

We consider a population genetics model of UD gene drive in a panmictic (randomly mating), isolated (closed) population of infinite size. Mutation is allowed to occur in each transgenic construct at a constant rate (*m* per gene) and we assume that *m* is low enough for multiple mutations in a single generation to be neglected. This gives a total of 18 alleles, with nine at each genetic locus (i.e. *a*, *A*, *A*_*L*_, *A*_*S*_, *A*_*C*_, *A*_*LS*_, *A*_*LC*_, *A*_*SC*_, *A*_*LSC*_, *b*, *B*, *B*_*L*_, *B*_*S*_, *B*_*C*_, *B*_*LS*_, *B*_*LC*_, *B*_*SC*_ and *B*_*LSC*_) where lower case denotes wild-type alleles, upper case represents transgenic alleles and subscripts indicate loss-of-function mutations in a given gene (*L*=lethal, *S*=suppressor and *C*=cargo). This results in a total of 2,025 possible genotypes, 819 of which are non-viable.

The fitness of each genotype is expressed relative to wild-type and represents a reduction in survival due to the carrying of transgenic constructs. Many genotypes also suffer a lethal effect from non-suppressed lethal genes. These factors are combined to give the relative fitness of each genotype:

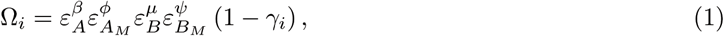

where *ε* denotes the relative fitness per construct conferred by non-mutated *(A, B)* or mutated (*A*_*M*_, *B*_*M*_ where *M* = *L, S, C, LS, LC, SC, LSC*) transgenic constructs. We assume relative fitnesses act multiplicatively and that all mutated transgenic constructs confer the same relative fitness regardless of the type or number of mutations (although resulting genotypes may confer a separate lethal effect). Exponents *β*, *ϕ*, *μ* and *ψ* denote the number of each transgenic construct type carried by a given genotype while *γ*_*i*_ represents the lethality of that genotype (*γ*_*i*_ = 1 if non-viable or *γ*_*i*_ = 0 if viable). Finally, we assume all lethal effectors are 100% effective when unsuppressed and that any number (i.e. one or two) of lethal effector copies are fully suppressed by any number (one or more) of the relevant suppressor (i.e. the strong suppression case in [9]).

**Figure 2:**
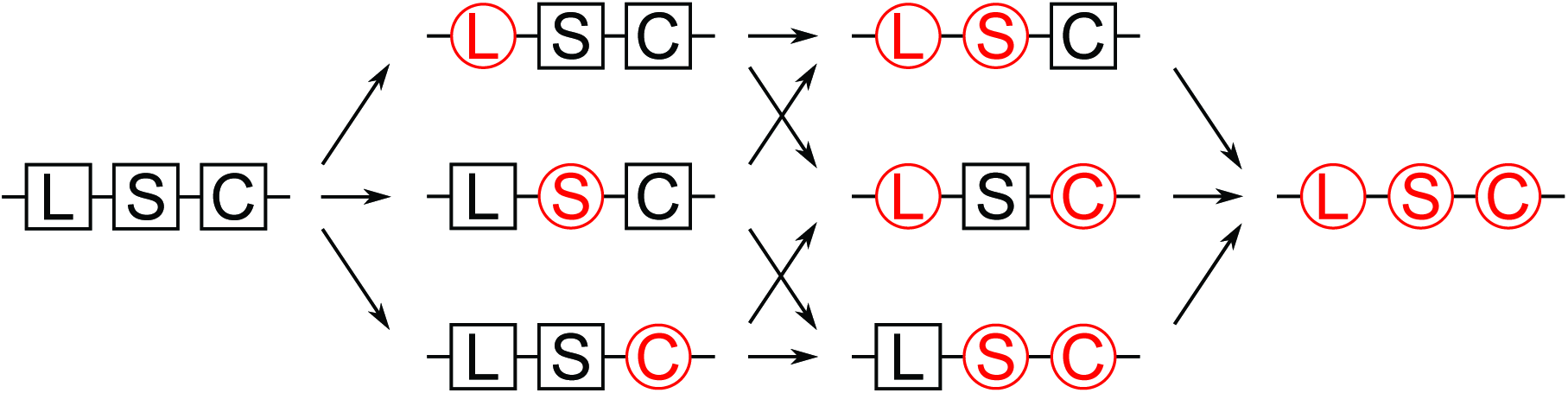
Transgenic constructs are assumed to mutate at a rate m per gene. This is assumed low enough that multiple mutations per generation may be neglected. For example the initial transgenic construct (say, *A*) mutates at a rate of 3*m*, producing mutations in the lethal (giving *A*_*L*_), suppressor(*A*_*S*_) and the cargo (*A*_*C*_) gene each at a rate of *m*. Then, transgenic constructs possessing one mutated gene (e.g. *A*_*L*_) mutate at a rate of 2*m* (giving *A*_*LS*_ and *A*_*LC*_ each at rate *m*). Transgenic constructs with two mutated genes (e.g. *A*_*LS*_) then mutate at rate *m* producing constructs with all three genes mutated (i.e. *A*_*LSC*_). Here non-mutated genes are represented by black squares whereas genes with loss-of-function mutations are shown in red circles.

For numerical simulations we used a set of MATLAB (MATLAB R2014b, The MathWorks Inc., Natick, MA) scripts run in parallel using the MATLAB Parallel Computing Toolbox through a MATLAB Distributed Computing Server. These allow simulation of the system without a manual formulation of 2,025 difference equations (see Figure 3). Briefly, the process begins by converting each possible genotype into a numerical form and computing relative fitnesses for each (using equation (1)). Initial conditions are then calculated according to:

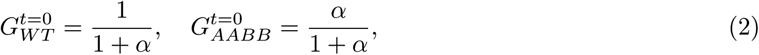

where 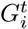 is the *i*-th genotype frequency *t* generations after the initial release; *α* is the release ratio (introduced/wild) of non-mutated transgene double homozygotes *(AABB)* into a wild-type *(aabb)* population; and all other genotypes have an initial frequency of zero (i.e. 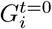 = 0 for *i ≠ WT, AABB*). We assume the absence of mutations in released individuals is feasible due to quality controls at the rearing facility. Genotype frequencies in subsequent generations are computed iteratively. Expected (Mendelian) frequencies are first calculated using a matrix of outcomes from every genotype mating pair and then multiplied by a matrix of mutation rates (giving proportional frequencies 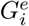). Finally, proportional genotype frequencies (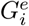) are normalized using the average fitness of the population (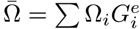) such that genotype frequencies (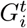) sum to one.

**Figure 3:**
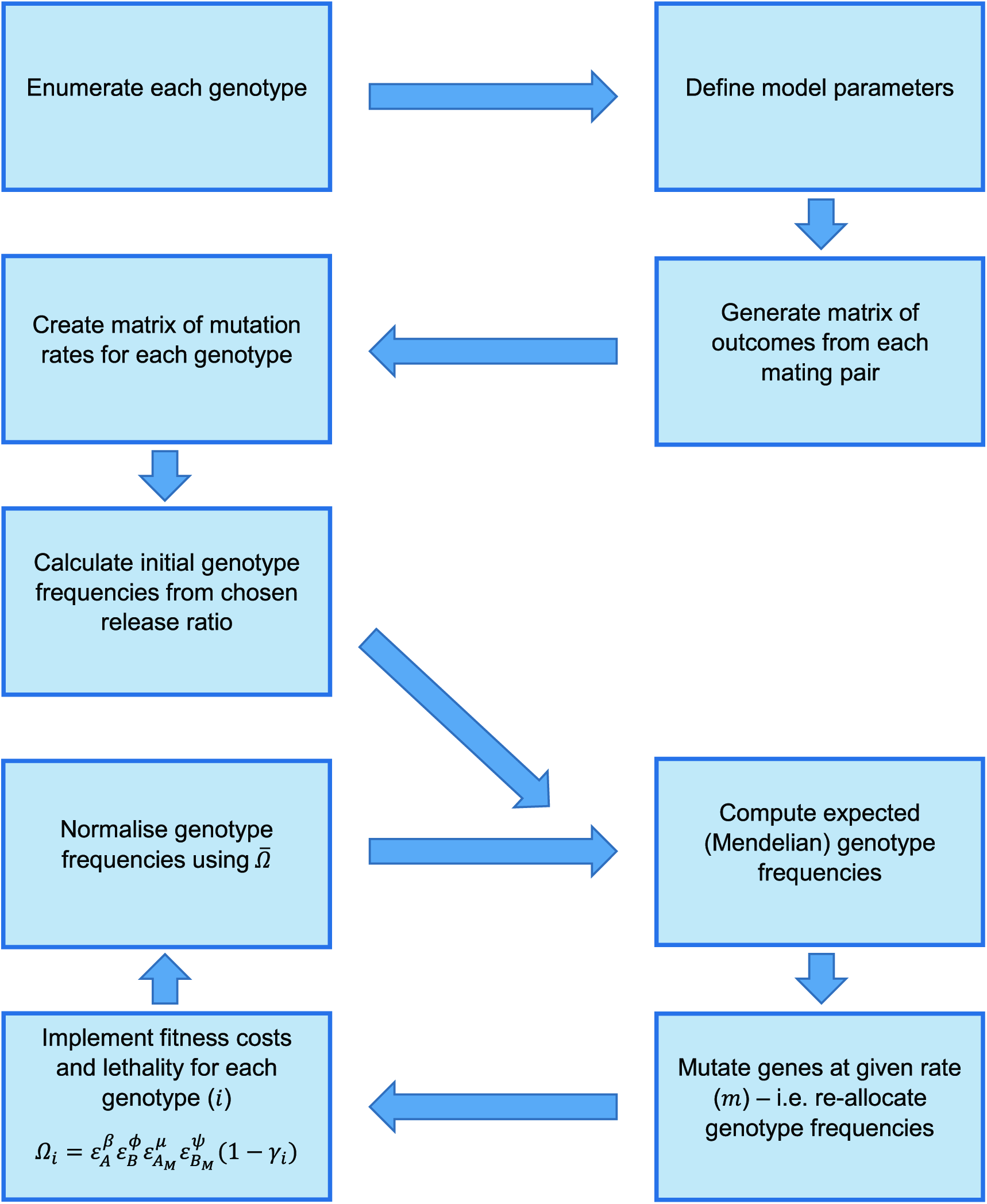
Diagram showing the simulation procedure for an engineered underdominance system with a constant rate of mutation (m) per gene. Parameter symbols used here are: *ε*, the relative fitness conferred by a given allele (denoted in the subscripts where M represents a mutated allele); *β* and *ϕ*, the numbers of non-mutated copies of construct A and B, respectively; *μ* and *ψ*, the number of mutated copies of transgenic constructs A and B, respectively; *γ*_*i*_, the lethality conferred by a given genotype (*i*); *Ωi*, the overall relative fitness of individuals of genotype *i*; and 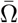, the average relative fitness of the entire population.

**Table 1:** Table of parameter and variable definitions used throughout this study.

| Symbol | Definition |
| --- | --- |
| $G_i$ | Frequency of genotype ($i$) |
| $G_i^e$ | Proportional (i.e. non-normalized) genotype frequency |
| $m$ | Rate of mutation per gene |
| $\alpha$ | Release ratio ($\alpha$=introduced/wild) |
| $\bar{\Omega}$ | Average fitness of the overall population |
| $\Omega_i$ | Relative fitness of genotype ($i$) |
| $\varepsilon_j$ | Relative fitness conferred by transgenic construct ($j$) |
| $\beta$ | Number of non-mutated copies of transgenic construct A |
| $\phi$ | Number of mutated copies of transgenic construct A (i.e. $A_M$) |
| $\mu$ | Number of non-mutated copies of transgenic construct B |
| $\psi$ | Number of mutated copies of transgenic construct B (i.e. $B_M$) |
| $\gamma_i$ | Lethality of genotype ($i$) |

## 3 Results

### 3.1 Numerical simulations with various rates of mutation per gene

To our knowledge, experimentally measured rates of mutation in key organisms considered as likely targets for UD gene drives are not well known. Thus, here we conduct numerical simulations for rates of mutation (per gene) spanning five orders of magnitude, namely *m* = 10^−4^, 10^−5^, 10^−6^, 10^−7^ and 10^−8^. Given previous estimates of the mutation rate in *Drosophila melanogaster* (2.8×10^−9^ [14] and 8.4×10^−9^ [15] per nucleotide per generation), the size of gene drive components used in previous studies (1-10kb e.g. [16-18]) and an approximation that 1-10% of nucleotides in these sequences are essential (i.e. causes a loss of gene function when mutated) we anticipate that this range should include rates relevant to a range of proposed target species.

Figure 4 shows results for 1:1 (introduced:wild) releases of double homozygote (i.e. *AABB*) individuals. We assume each non-mutated transgenic construct confers a fitness load of either 5% or 10% (i.e. *ε*_*A*_ = *ε*_*B*_=0.95 or 0.90) and mutated constructs 4% or 8% (i.e. *ε*_*A*_*M*__ = *ε*_*B*_*M*__=0.96 or 0.92), respectively. Thus, mutated transgenic constructs have a small fitness advantage over non-mutated ones, yet still a deficit relative to wild-type. In examples conducted with mutated transgenic alleles conferring a greater fitness cost than non-mutated alleles, the UD system progressed without significant accumulation of mutated alleles, thus we do not consider these cases any further.

**Figure 4:**
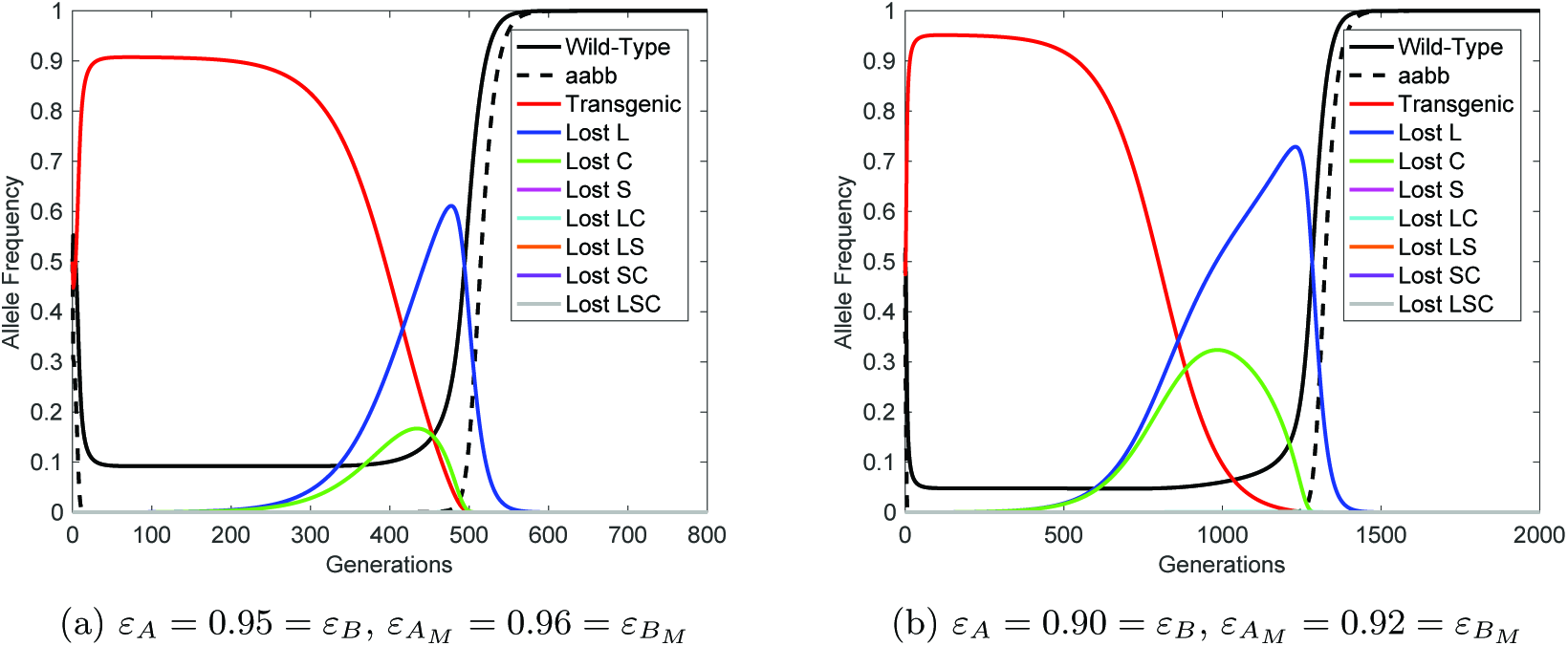
Mutation of transgenic constructs is predicted to return a fully wild-type population. Results here are presented for 1:1 (introduced:wild) introductions of non-mutated double homozygote *(AABB)* individuals into a wild-type *(aabb)* population. Here, transgenic constructs are assumed to mutate at a rate *m* = 10^−6^ per gene, neglecting the possibility of multiple genes mutating within a single generation. Solid lines represent the allele frequency of wild-type, transgenic and each variety of mutated allele with details given in the figure legend whilst black dashed lines denote the genotype frequency of double wild-type homozygotes. Here a number of alleles only reach very low maximum frequencies (see Figure 5(a)) and thus they appear to overlie one another along the horizontal axis. Panel (a) shows results for *ε*_*A*_ = 0.95 = *ε*_*B*_ and *ε*_*A*_*M*__ = 0.96 = *ε*_*B*_*M*__ whereas (b) is for *ε*_*A*_ = 0.90 = *ε*_*B*_ and *ε*_*A*_*M*__ = 0.92 = *ε*_*B*_*M*__. Note the difference in time-scales between these panels.

In Figure 4 the non-mutated UD system initially reaches a high frequency, as previously modeled [5-11]. However, each type of mutated construct begins to accumulate in the population, with large frequencies reached by constructs carrying a single mutation in either the lethal or cargo gene. This results in a concurrent decrease in the frequency of non-mutated constructs. Once these mutated constructs have reached a high frequency in the population, they begin to be replaced by the remaining wild-type alleles due to the relative fitness advantage of wild-type alleles over mutated transgenic alleles. This eventually returns the population to a fully wild-type state.

For the full range of mutation rates considered here we observe similar dynamics to those in Figure 4 except that increasing (decreasing) mutation rates lead to faster (slower) accumulation and higher (lower) maximum frequencies of most types of mutated transgene allele (see Figure 5(a)). This inevitably means that higher rates of mutation reduce the period over which the UD system persists (see Figure 5(b)) and also the period for which functional cargo genes are present at high frequency (see Figure 5(c)). This should form a reasonable proxy for the efficacious period of a released UD system.

**Figure 5:**
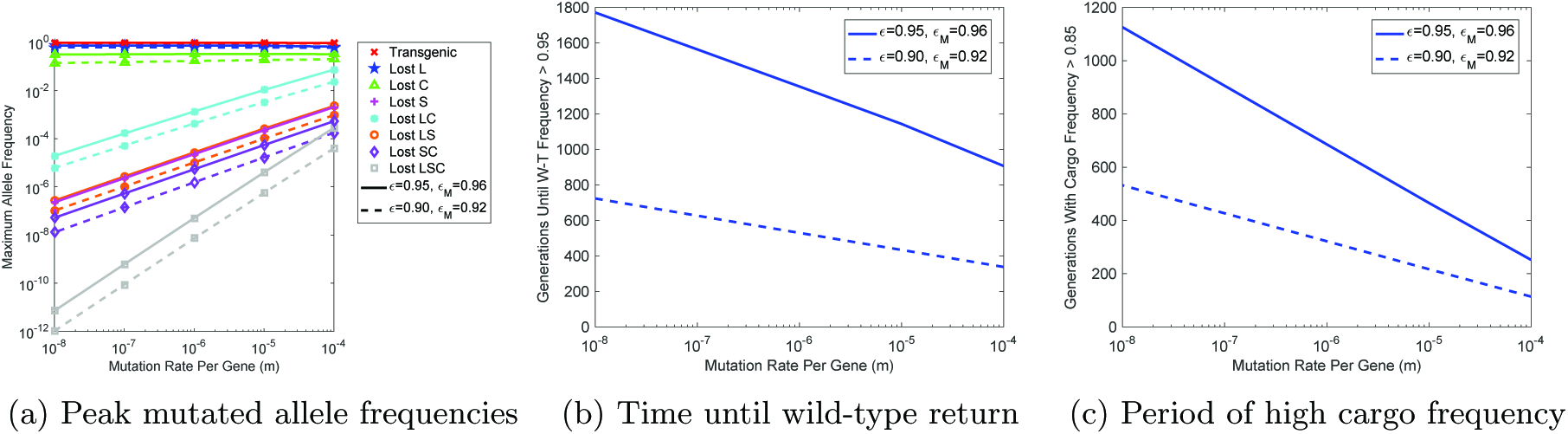
Rates of mutation in introduced engineered underdominance (UD) transgenes affect mutated allele frequencies and the efficacious period of the system. (a) Maximum frequencies attained by each of the non-mutated and mutated transgene alleles. Colors represent each different type of allele with details given in the figure legend. (b) The number of generations from the time of initial transgenic release until wild-type alleles return to a frequency greater than 0.95. (c) The number of generations that the UD system is able maintain a high frequency (>0.85) of transgenes with a functional copy of the cargo gene. In all panels solid lines represent cases with relative fitness parameters *ε*_*A*_ = 0.95 = *ε*_*B*_ and *ε*_*A*_*M*__ = 0.96 = *ε*_*B*_*M*__ whilst dashed lines are for cases with *ε*_*A*_ = 0.90 = *ε*_*B*_ and *ε*_*A*_*M*__ = 0.92 = *ε*_*B*_*M*__. These timings should represent a reasonable proxy for the period over which the desired phenotype conferred by the UD system would be effective.

### 3.2 Reversal strategies

To our knowledge, the only reversal strategy previously proposed for UD systems is the release of wild-type individuals in sufficient numbers to push the transgene frequency to sub-threshold levels [12, 13]. When successful, wild-type alleles should recover and eliminate transgenes from the population. Whilst this mechanism has been mentioned a number of times previously it has yet to be explored in any detail.

Based on results presented in Figure 4 here we propose an alternative, genetics-based, reversal strategy for UD systems. In this strategy transgenic individuals carrying only the suppressor genes of the original UD system would be released. These “free suppressor” carrying individuals are assumed to be of greater fitness than those carrying the original UD constructs since they carry less genetic material and will also suffer no lethal effect.

We conduct numerical simulations in order to compare the different reversal strategies discussed here. Similar to the examples above, we consider the relative fitness conferred by the initial UD system to be *ε*_*A*_ = 0.95 = *ε*_*B*_ or *ε*_*A*_ = 0.90 = *ε*_*B*_ and the free suppressor elements to confer a relative fitness of *ε*_Rev_ = 0.96 or *ε*_Rev_ = 0.92, giving them a small fitness advantage over the original UD constructs and a deficit relative to wild-type. Results are shown in Figure 6 for a 2:1 (introduced:wild (all genotypes)) release of wild-type individuals and individuals carrying free suppressors at both loci. These releases are assumed to occur 100 generations after an initial 1:1 (introduced:wild) UD release so that the original system is at high frequency and mutations would not have accumulated to significant frequencies. As such we initially assume that no mutation occurs (i.e. *m* = 0) to ensure that free suppressors work in absence of other types of allele.

**Figure 6:**
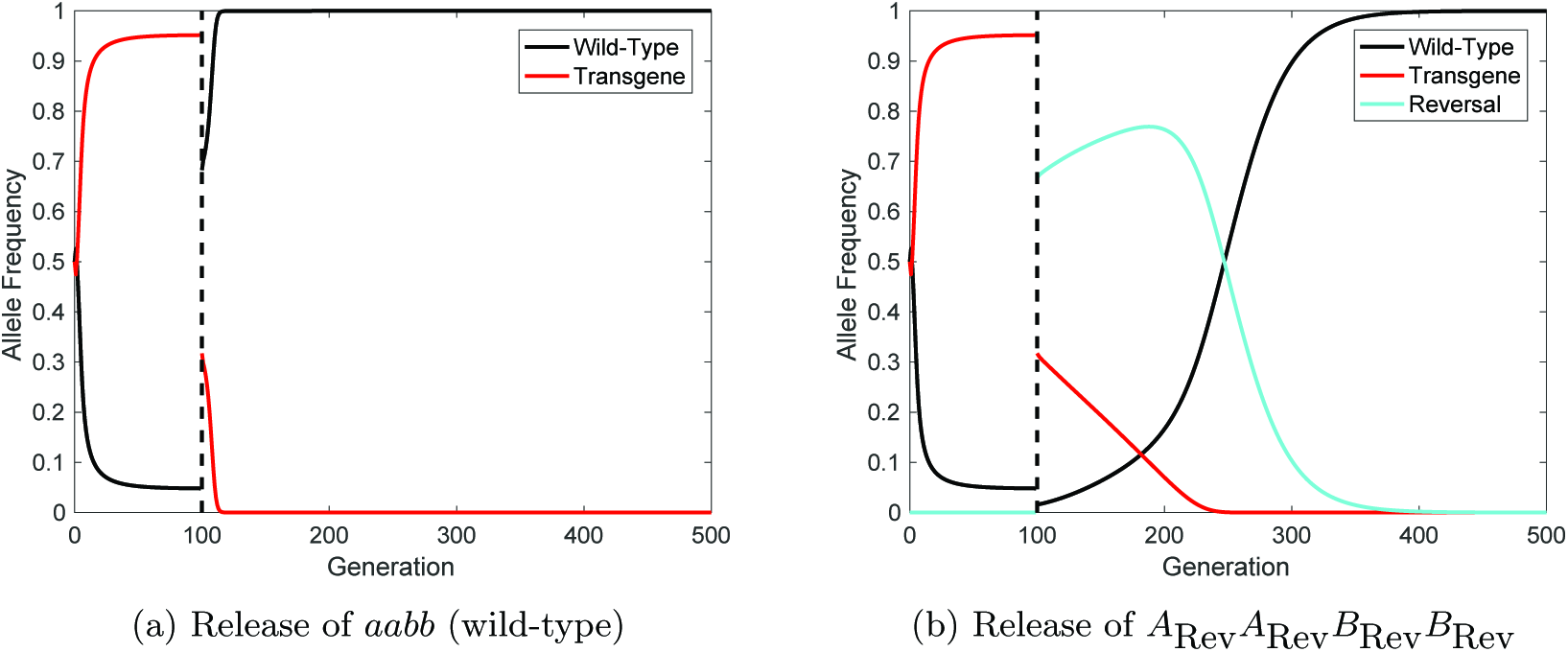
Comparison of two different reversal strategies for engineered underdominance (UD) gene drive systems. (a) An example of reversal through the release of wild-type *(aabb)* individuals. (b) Release of individuals carrying free suppressors at both loci (i.e. A _Rev_ A _Rev_ B_Rev_ B_Rev_). Here line colors denote each type of allele with black representing wild-type (a, b); red denoting non-mutated transgenes *(A, B);* and cyan showing suppressor-only alleles (*A*_Rev_, *B*_Rev_). In each example the initial UD release is made at a 1:1 (introduced:wild) ratio with relative fitness parameters *ε*_*A*_ = 0.95 = *ε*_*B*_. After 100 generations one of the reversal strategies is released at a ratio of 2:1 (introduced:wild).

Figure 6(b) demonstrates the feasibility of reversing of UD systems via the introduction of free suppressors in the absence of mutated alleles. We then relax this assumption to consider the behavior of a reversal drive in the presence of mutation (here we assume *A*_Rev_ = *A*_*LC*_ and *B*_Rev_ = *B_LC_*). These numerical simulations demonstrate that this reversal strategy can function even with small releases (i.e. *α* = 0.01 and *α* = 0.1; see Figure 7). However, one would need to be wary of stochastic effects when making extremely small releases. These results imply that the release of free suppressor carrying individuals could represent a very cost effective reversal strategy for UD systems.

**Figure 7:**
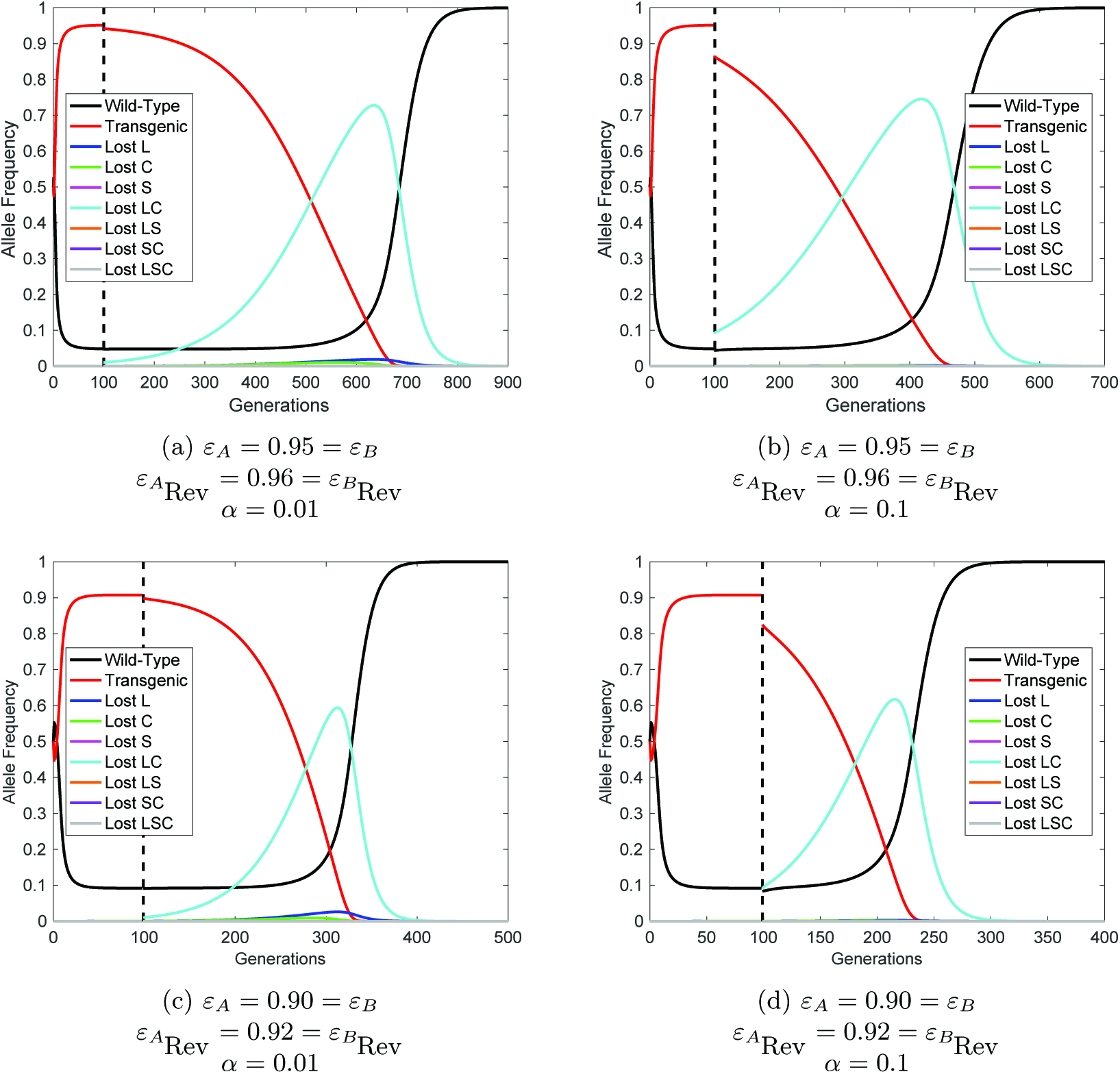
Free suppressor constructs can reverse engineered underdominance (UD) gene drives with small releases. Here, releases at generation 100 of free suppressor constructs (assuming *A*_Rev_ = *A*_*LC*_ and *B*_Rev_ = *B*_*LC*_) are numerically simulated for release ratios of *α* = 0.01 (panels (a) and (c)) and *α* = 0.1 (panels (b) and (d)) - far smaller than those required for reversal with wild-type. Lines represent the possible alleles with colors denoting the nature of each (details given in figure legends).

It is clear that release of wild-type individuals provides a much faster option for returning a fully wild-type population (Figure 6). However, releases of wild-type individuals must be significantly larger (∼1.75 times the wild population when *ε*_*A*_ = 0.95 = *ε*_*B*_)in order to achieve reversal of the initial UD system. This likely makes the release of wild-type individuals a far more costly reversal strategy than the use of free-suppressor carrying individuals.

## 4 Discussion

Since gene drive systems were first proposed they have gained much attention for their potential to help fight a number of important global issues (e.g. [1,2,19,20]). However, various genetic control measures have been shown to be limited by the potential generation of mutation or resistance [18,21-25]. A number of studies have also begun to investigate mechanisms capable of reversing gene drives in case they produce unexpected consequences [26-28]. Here we showed that mutations will likely lead to the break down of UD systems and the eventual elimination of introduced transgenes although under the assumptions used, the UD system is likely to persist at high allele frequency for hundreds of generations. We then went on to demonstrate that reversal of UD systems is feasible via the release of either wild-type individuals or individuals carrying free suppressors.

As with all mathematical models, the work presented here relies on a number of simplifying assumptions, the majority of which are common in this type of study [5,6,9,11]. Since these assumptions have been discussed previously, we do not consider them any further here. There are however a few areas specific to this study where more detailed modeling would be useful to further elucidate the effects of mutation in UD systems. Firstly, we assumed that mutations completely eliminate gene function whereas in reality they may produce only a partial loss of function. We also assumed that all mutated transgenic constructs confer the same fitness cost regardless of the number and type of mutations carried. These assumptions could be relaxed in future models by expressing gene efficacy as a function of mutations carried. Finally, it is possible that natural genetic polymorphism in a target population could mean that certain individuals would be, at least partially, resistant to the effects of introduced genes. In such a case, the UD system may place resistance alleles under a selective pressure causing them to increase in frequency and reduce the efficacy of UD systems.

Results presented here assume that individuals carrying mutated transgenes are fitter than those carrying non-mutated transgenic constructs. This is a common assumption when modeling the formation of resistance/mutations in gene drive systems (e.g. [21, 24]). The fitness advantage of individuals carrying specific mutations in each transgene (or combination of transgenes) is likely to determine the exact dynam-ics and time scales that would be observed in experimental work. Results presented here are intended to give a guide as to the type of behavior we would expect to emerge rather than giving definitive predictions for specific applications.

Previous literature has estimated the mutation rate in Drosophila melanogaster to be ∼ 5.6×10^−9^ per nucleotide per generation (mean of estimates in [14] and [15]). Given the size of gene drive components used in previous experimental studies (1-10kb e.g. [16-18]) and an approximation that 1-10% of nucleotides in these sequences are essential (i.e. causes a loss of gene function when mutated), we believe the mutation rates explored here to be feasible for a range of proposed target organisms. Using these mutation rates, we assumed only one mutation was able to emerge in a single generation. In practice, deletions could remove two adjacent components simultaneously. However, assuming the ordering of components shown in Figure 1 such deletions would leave individuals without suppressor elements, thus making them non-viable in presence the of the construct at the other locus. Coupling this with the extremely low probability of multiple mutations occurring simultaneously, we do not anticipate that relaxing this assumption would lead to major differences from the results presented here.

A number of recent studies have discussed the need to explore reversal strategies for gene drives in case they have unexpected consequences. While the majority of work has focused on the reversal of CRISPR-Cas9 gene drive systems (e.g. [26-29]), it has been suggested that the release of wild-type individuals could reverse UD systems [12,13]. The release of wild-type individuals represents a threshold dependent reversal strategy with the precise number of wild-types required to reverse a released UD system depending on the fitness of UD carrying individuals and the size of the wild population, both of which are difficult to measure. What is known however, is that the size of these wild-type releases would need to be very large. For example, when transgenic constructs each confer a 5% fitness cost (i.e. *ε*_*A*_ = 0.95 = *ε*_*B*_), the wild-type release would need to be ∼1.75 times the size of the entire wild population. Importantly, insufficiently sized releases of wild-type individuals would fail to reverse a given UD system. While it would be feasible to make further wild-type releases, this would have obvious cost implications. To help overcome these issues, here we proposed a genetics-based alternative (release of free suppressor carriers) that appears to be threshold independent (i.e. it is predicted to be effective from very small releases, leaving aside stochastic effects) but will take longer to return a fully wild-type population than release of wild-types. It may also be difficult to convince the public that releasing further transgenics is an acceptable method of eliminating a gene drive that produced unexpected consequences. Here we do not wish to draw a firm conclusion on which reversal strategy should be pursued, since this may depend on case-specific social, economic and technical factors that are beyond the scope of this study.

In spite of demonstrating that UD gene drives will likely break down over time with the emergence of mutations, this study provides reason to be optimistic about the prospect of using UD gene drives to spread desirable genes through a target population. In particular, even though introduced transgenes are likely to be eliminated, the desirable genetic cargo is expected to reach and maintain a high frequency in the population for hundreds of generations. This is likely long enough for them to have produced their desired effect. Here we have given the first theoretical examination of this UD break down due to mutation. We anticipate that future modeling studies will be able to produce more application specific models, thus refining the predictions presented here.

## Acknowledgements

MPE is funded through a Wellcome Trust Investigator Award [110117/Z/15/Z] made to LSA. LSA is supported by core funding from the UK Biotechnology and Biological Sciences Research Council (BBSRC) to The Pirbright Institute [BBS/E/I/00007033].

